# Genetic differentiation of *Xylella fastidiosa* following the introduction into Taiwan

**DOI:** 10.1101/2021.03.24.436723

**Authors:** Andreina I. Castillo, Chi-Wei Tsai, Chiou-Chu Su, Ling-Wei Weng, Yu-Chen Lin, Shu-Ting Cho, Rodrigo P. P. Almeida, Chih-Horng Kuo

## Abstract

The economically important plant pathogen *Xylella fastidiosa* has been reported in multiple regions of the globe during the last two decades, threatening a growing list of crops and industries. *Xylella fastidiosa* subspecies *fastidiosa* causes disease in grapevines (Pierce’s disease of grapevines, PD), a current problem in the United States (US), Spain, and Taiwan. We studied PD-causing subsp. *fastidiosa* populations and compared the genome sequences of 33 isolates found in Central Taiwan with 171 isolates from the US and two from Spain.

Phylogenetic relationships, haplotype network, and genetic diversity analyses confirm that subsp. *fastidiosa* was recently introduced into Taiwan from the Southeast US (i.e., the PD-I lineage in Georgia based on available data). Recent core genome recombination events were detected among introduced subsp. *fastidiosa* isolates in Taiwan and contributed to the development of genetic diversity, particularly in the Houli District of Taichung City in Central Taiwan. Unexpectedly, despite comprehensive sampling of all regions with high PD incidences in Taiwan, the genetic diversity observed include contributions through recombination from unknown donors, suggesting that higher diversity exists in the region. Nevertheless, no recombination event was detected between *X. fastidiosa* subsp. *fastidiosa* and the endemic sister species *Xylella taiwanensis*. In summary, this study improved our understanding of the genetic diversity of PD-causing subsp. *fastidiosa* after invasion to a new region.

## Introduction

Southeast Asia is a region of intricate tectonic and climatic evolution, complex biogeography, and a hotspot for global biodiversity (de Bruyn et al. 2014; Hughes 2017; Lohman et al. 2011; Sholihah et al. 2021). Oceanic islands like Taiwan, are important contributors to plant richness within the region, with local diversity reflecting both palaeogeographic events and human mediated plant movement (Liao and Chen 2017). Though there are many species endemic to Taiwan, approximately 73% of vascular plants found in the island originated from neighboring regions (Hsieh 2002). In recent decades, a significant number of plants from the Americas, Europe, and Africa have also been introduced. By the start of the new millennium, it is estimated that 341 plant species have been naturalized in Taiwan (Wu et al. 2004), many of them crops. In association with these novel crop species, numerous pests and pathogens have also been introduced (Yeh 2005; Wu 2006).

The economic losses caused by phytopathogen infections are estimated as $1 billion dollars worldwide every year (Martins et al. 2018). In addition to the economic costs, emerging plant pathogens represent a threat to food security, not only by affecting a region’s capacity to meet its nutritional needs, but also by negatively impacting local economies and social structures dependent on trade (Fones et al. 2020). The difficulties associated with the control and management of introduced plant diseases are well documented (He et al. 2016). Specifically, the genetic homogeneity of crop monocultures favors the emergence of host-specialized pathogen genotypes and the increment of virulence (McDonald and Stukenbrock 2016; Read 1994). Furthermore, pathogens can quickly adapt to geographical and ecological conditions (Croll and McDonald 2017; Dutta et al. 2021; Giraud et al. 2010; Mhedbi-Hajri et al. 2013). These problems are particularly concerning in the context of introduced plant pathogens with wide host ranges and demonstrated capacity to spill over to sympatric flora. In this regard, the xylem-limited bacterial plant pathogen *Xylella fastidiosa*, transmitted by various insect vectors, represents an important topic of study (reviewed by Sicard et al. 2018). The bacterium *X. fastidiosa* is taxonomically divided into three main monophyletic subspecies with allopatric origins: *X. fastidiosa* subsp. *fastidiosa*, native to Central America (Castillo et al. 2020; Nunney et al. 2019; Vanhove et al. 2020); *X. fastidiosa* subsp. *multiplex*, native to temperate and subtropical North America (Nunney et al. 2012, 2014); and *X. fastidiosa* subsp. *pauca*, thought to be native to South America (Nunney et al. 2012). Each of these subspecies has experienced a recent geographical expansion mediated by unique intra- and inter-continental introduction events (Gomila et al. 2018; Landa et al. 2020; Olmo et al. 2017; Saponari et al. 2018; Sicard et al. 2018; Vanhove et al. 2020). In the particular case of subsp. *fastidiosa*, that includes one introduction to the United States (US) that led to the emergence of Pierce’s disease (PD) (Vanhove et al. 2020) and a subsequent introduction to the island of Mallorca in Spain (Gomila et al. 2018). In addition, the invasion of subsp. *fastidiosa* in Central Taiwan dates back to at least 2002, when the pathogen was first isolated from *Vitis vinifera* L. plants showing symptoms of PD (Su et al. 2013a). Later studies suggested that this invasion originated via the introduction of PD-causing subsp. *fastidiosa* from the Southeast United States (US), also known as the PD-I lineage (Castillo et al. 2019, 2021). Local xylem feeders (*Kolla paulula* and *Bothrogonia ferruginea*) capable of transmitting subsp. *Fastidiosa* have been identified (Lin and Chang 2012; Tuan et al. 2016). As a consequence, PD control strategies have mainly been centered around vector control, assessment of the habitats suitable for these vectors, eradication of infected plants and alternative host plants, and planting of healthy seedlings (Su et al. 2013b). There is another xylem-limited pathogen only found in Taiwan. Leaf scorch symptoms described in pear plants (*Pyrus pyrifolia*) in Central Taiwan were documented in the 1980s, and the appearance of disease symptoms was correlated with the presence of a xylem-limited fastidious bacterium (Leu and Su 1993). Isolate PLS229 was obtained from symptomatic pear cultivars “Hengshan” and sequencing of its 16S rRNA showed 99% similarity with published *X. fastidiosa* isolates (Su et al. 2014). Subsequent analyses found that nucleotide identity (ANI) between PLS229 and *X. fastidiosa* ranged between 83.4-83.9%, suggesting that PLS229 was a new species (Su et al. 2016). The species, named *Xylella taiwanensis*, represents the only currently known sister taxon to *X. fastidiosa*. Analysis of the complete genome of *X. taiwanensis* shows that this species shares only ∼66-70% of its gene content with *X. fastidiosa* (Weng et al. 2021). Both *X. taiwanensis* and *X. fastidiosa* subsp. *fastidiosa* have a negative effect on Taiwanese crops. In the case of subsp. *fastidiosa*, vineyards located in hilly terrains are under high risk of PD infection. In an analysis of ten-year survey data, it was shown that infection rate of grapevines was highest in the Waipu District of Taichung City (69%), Tonxiao Township of Miaoli County (87%), and Houli District of Taichung City (93%) (Su et al. 2013b). Nonetheless, studies are yet to assess the genetic diversity in subsp. *fastidiosa* isolates found within these regions or establish the degree of local-adaptation and genotypic divergence among areas. Moreover, the prevalence of evolutionary events such as gene gain/loss, homologous recombination, and mutation rate remain to be established. Addressing these points is important to understanding how subsp. *fastidiosa* adapts to new areas following an introduction event, and it is also crucial in aiding the implementation of disease management strategies in Taiwan.

We conducted the first analysis of the genomic diversity of PD-causing subsp. *fastidiosa* isolates found in grapevines collected from Central Taiwan, covering most of the known geographical range in the area. We examined the ancestor/descendant relationships among introduced PD-causing subsp. *fastidiosa* populations. We also evaluated trends of genetic diversity within Taiwan and contrasted them to other worldwide PD-causing subsp. *fastidiosa* populations. Finally, we assessed the role that homologous recombination has in the development of genetic diversity across regions in Taiwan.

## Materials and Methods

### Strain isolation and genome sequencing

Thirty-one field isolates were collected from symptomatic grapevines in Central Taiwan (Table 1). Xylem-limited bacteria were isolated as previously described (Su et al. 2013a). Most isolates were collected from the Waipu and Houli Districts in Taichung City, and two isolates were from Tonxiao and Zhuolan Townships in Miaoli County, respectively (Figure 1b). The procedures for whole genome shotgun sequencing and data processing were based on those described in our previous studies (Castillo et al. 2021; Weng et al. 2021). All bioinformatics tools were used with the default settings unless stated otherwise.

**Table 1.** Summary of assembly statistics for the newly sequenced Taiwanese *Xylella fastidiosa* subsp. *fastidiosa* genomes

| Isolate | Geographic origin | Grapevine variety | Collection date | N50 (bp) | Genome size (bp) | Coverage (x) |
| --- | --- | --- | --- | --- | --- | --- |
| GV210 | Tongxiao, Miaoli | Kyoho | 2015.08.21 | 87,160 | 2,463,940 | 294 |
| GV215 | Houli, Taichung | Kyoho | 2017.06.30 | 99,605 | 2,463,006 | 331 |
| GV216 | Houli, Taichung | Kyoho | 2017.08.03 | 97,514 | 2,468,540 | 285 |
| GV219 | Waipu, Taichung | Kyoho | 2017.07.24 | 92,425 | 2,460,764 | 237 |
| GV220 | Waipu, Taichung | Kyoho | 2017.07.24 | 99,605 | 2,461,512 | 288 |
| GV221 | Waipu, Taichung | Kyoho | 2017.07.24 | 94,057 | 2,461,485 | 258 |
| GV222 | Waipu, Taichung | Kyoho | 2017.07.24 | 113,299 | 2,461,579 | 254 |
| GV225 | Waipu, Taichung | Kyoho | 2017.07.24 | 92,981 | 2,470,584 | 272 |
| GV229 | Waipu, Taichung | Golden Muscat | 2017.07.24 | 99,551 | 2,465,605 | 232 |
| GV230* | Waipu, Taichung | Black Queen | 2018.06.24 | 2,514,993 | 2,514,993 | 277 |
| GV231 | Waipu, Taichung | Golden Muscat | 2018.06.24 | 101,467 | 2,490,832 | 274 |
| GV232 | Waipu, Taichung | Golden Muscat | 2018.06.24 | 90,280 | 2,467,990 | 272 |
| GV233 | Waipu, Taichung | Golden Muscat | 2018.06.24 | 100,549 | 2,461,948 | 209 |
| GV234 | Waipu, Taichung | Golden Muscat | 2018.06.24 | 100,555 | 2,463,675 | 213 |
| GV235 | Waipu, Taichung | Golden Muscat | 2018.06.24 | 100,555 | 2,466,447 | 197 |
| GV236 | Waipu, Taichung | Golden Muscat | 2018.06.24 | 100,555 | 2,465,286 | 215 |
| GV237 | Waipu, Taichung | Golden Muscat | 2018.06.24 | 92,561 | 2,468,425 | 207 |
| GV238 | Waipu, Taichung | Golden Muscat | 2018.06.24 | 90,280 | 2,516,844 | 285 |
| GV239 | Waipu, Taichung | Golden Muscat | 2018.06.24 | 99,551 | 2,490,161 | 250 |
| GV240 | Waipu, Taichung | Golden Muscat | 2018.06.24 | 124,734 | 2,469,727 | 237 |
| GV241 | Waipu, Taichung | Golden Muscat | 2018.06.24 | 90,280 | 2,467,728 | 219 |
| GV244 | Waipu, Taichung | Golden Muscat | 2018.12.25 | 80,059 | 2,774,440 | 57 |
| GV245 | Waipu, Taichung | Golden Muscat | 2018.12.25 | 92,729 | 2,600,059 | 83 |
| GV248 | Waipu, Taichung | Black Queen | 2019.07.03 | 98,380 | 2,456,198 | 19 |
| GV249 | Waipu, Taichung | Black Queen | 2019.07.03 | 86,393 | 2,456,603 | 15 |
| GV252 | Waipu, Taichung | Golden Muscat | 2019.07.03 | 90,026 | 2,455,757 | 16 |
| GV253 | Waipu, Taichung | Golden Muscat | 2019.07.03 | 99,557 | 2,458,547 | 15 |
| GV263 | Waipu, Taichung | Black Queen | 2019.12.04 | 97,977 | 2,460,836 | 14 |
| GV264 | Waipu, Taichung | Black Queen | 2019.12.04 | 95,089 | 2,461,237 | 17 |
| GV265 | Waipu, Taichung | Black Queen | 2019.12.04 | 99,087 | 2,457,696 | 16 |
| GV266 | Zhuolan, Miaoli | Kyoho | 2019.12.31 | 86,635 | 2,459,085 | 16 |

\* The reference isolate GV230 that has the complete genome sequence available.

**Figure 1.**
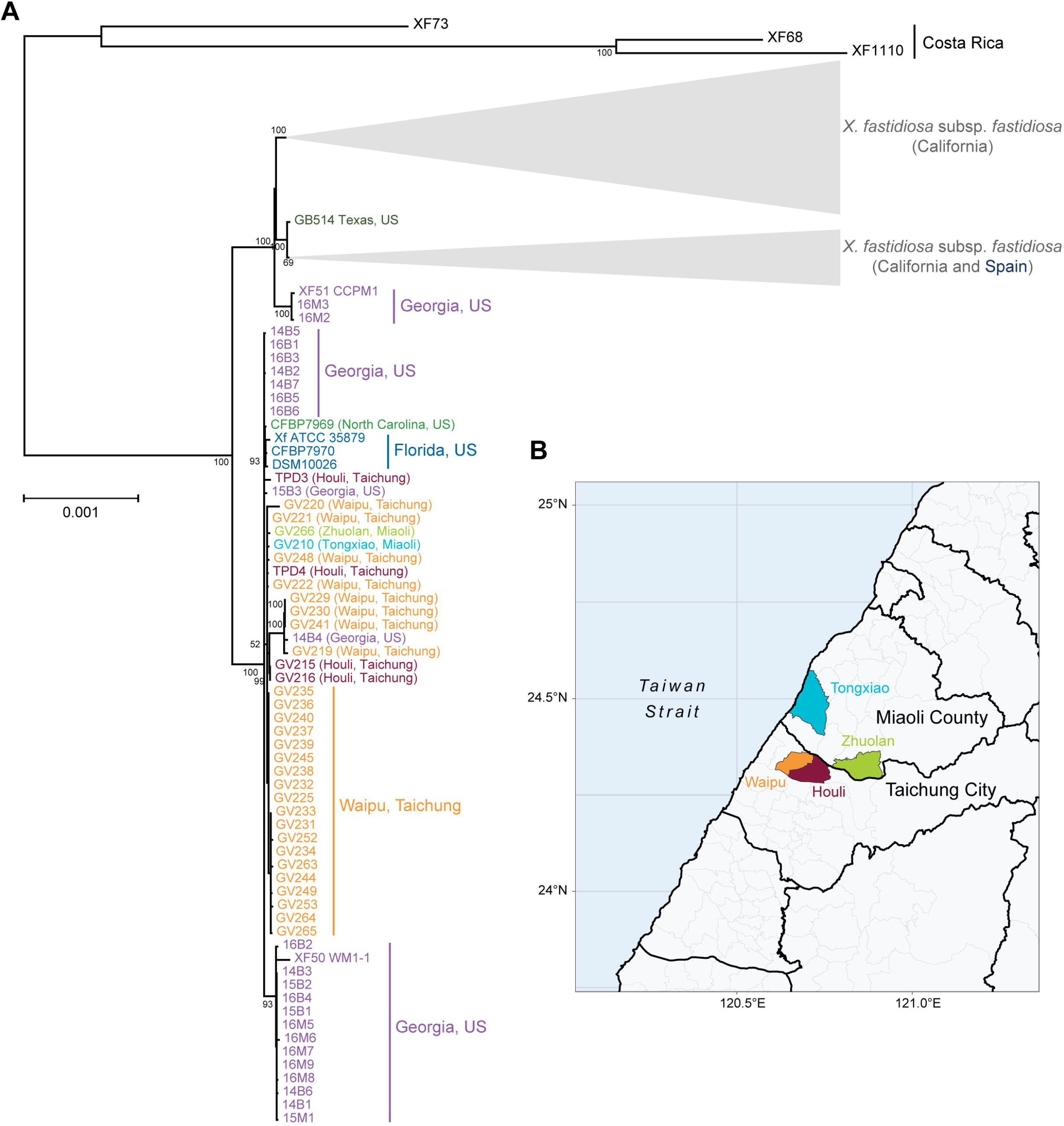
Relationship of PD-causing isolates. **a**. Maximum Likelihood (ML) tree of worldwide PD-causing subsp. *fastidiosa* isolates. The phylogenetic inference was based on the core genome (i.e., shared by > 99% of the isolates) without removal of recombinant segments. Costa Rican isolates were used to root the tree. Clades encompassing California and Spain isolates have been compressed to their most recent common ancestor. **b**. Collection locations of the Taiwan isolates.

Briefly, the DNA samples were prepared using the Wizard Genomic DNA Purification Kit (A1120; Promega, USA). The Illumina sequencing libraries were prepared by the Genomic Technology Core (Institute of Plant and Microbial Biology, Academia Sinica) using KAPA LTP Library Preparation Kit (KK8232; Roche, Switzerland) and KAPA Dual-Indexed Adapter Kit (KK8722; Roche, Switzerland) with a target insert size of ∼550 bp. The Illumina MiSeq paired-end sequencing service was provided by the Genomics Core (Institute of Molecular Biology, Academia Sinica) using the MiSeq Reagent Nano Kit v2 (MS-103-1003; Illumina, USA). For the reference isolate GV230, additional sequencing was conducted using the Oxford Nanopore Technologies (ONT) MinION platform (FLO-MIN106; R9.4 chemistry and MinKNOW Core v3.6.0) (Oxford Nanopore Technologies, UK). The sequencing library was prepared using the ONT Ligation Kit (SQK-LSK109) without shearing or size selection. Guppy v3.4.5 was used for base calling.

To obtain the complete genome sequence of the reference isolate GV230, the Illumina and ONT reads were combined for *de novo* assembly by using Unicycler v0.4.8-beta (Wick et al. 2017). For validation, the Illumina and ONT raw reads were mapped to the assembly using BWA v0.7.12 (Li and Durbin 2009; Wang et al. 2012) and Minimap2 v2.15 (Li 2018), respectively. The mapping results were programmatically checked using SAMtools v1.2 (Li et al. 2009) and manually inspected using IGV v2.3.57 (Robinson et al. 2011). Gene prediction and annotation were performed using the NCBI prokaryotic genome annotation pipeline (Tatusova et al. 2016).

For the remaining isolates, draft genome assemblies were prepared using only Illumina reads. The quality of raw paired FASTQ reads was evaluated using FastQC (Wingett and Andrews 2018). Low quality reads and adapter sequences were removed from all paired raw reads using seqtk v1.2 (https://github.com/lh3/seqtk) and cutadapt v1.14 (Marcel 2011). After pre-processing, SPAdes v3.13 (Bankevich 2012; Nurk et al. 2013) was used for *de novo* assembly with the -*careful* parameter and a -*k* of 21, 33, 55, and 77. Contigs were reordered with Mauve’s contig mover function (Rissman et al. 2009) using the complete assembly of GV230 as a reference. Assembled and reordered genomes were then individually annotated using the Prokka pipeline (Seemann 2014).

### Core genome alignments, construction of ML trees, and recombination detection

Roary v3.11.2 (Page et al. 2015) was used to calculate the number of genes within each genome fragment (core, soft-core, shell, and cloud genome) and generate a core genome alignment for worldwide PD-causing subsp. *fastidiosa* strains (N = 206). Three Costa Rican isolates (non-grapevine infecting) were used as an outgroup (Castillo et al. 2020). The core genome alignment was used to build a Maximum Likelihood (ML) tree with RAxML (Stamatakis 2014). The GTRCAT substitution model was used on tree construction, while tree topology and branch support were assessed with 1000 bootstrap replicates. A location-specific core genome alignment was generated using only the Taiwan isolates, including two published (Castillo et al. 2019) and 31 newly reported in this work. This is subsequently referred to as the Taiwan-only dataset (N = 33). Likewise, a core genome alignment was generated for the Taiwan-only isolates plus their ancestor population (Southeast US or PD-I). This is subsequently referred to as the PD-I/Taiwan dataset (N = 61). The ancestor/descendant relationships among populations were established using the worldwide PD-causing subsp. *fastidiosa* ML tree (see Results) as well as based on previous studies (Castillo et al. 2021).

The core genome alignments for worldwide PD-causing isolates were used to estimate the location of recombinant events in fastGEAR with default parameters (Mostowy et al. 2017). fastGEAR can identify lineage-specific recombinant segments (ancestral) and strain-specific recombinant segments (recent). Recombinant regions were fully removed from the alignment to generate a non-recombinant core genome. The non-recombinant core was realignment with MAFFT with default parameters (Katoh and Standley 2013) and used to construct a ML non-recombinant tree as previously described. It should be noted that fastGEAR was designed to test recombination in individual gene alignments instead of core genome alignments; yet, a previous study found that fastGEAR was more conservative than other recombination detection methods such as ClonalFrameML (Vanhove et al. 2020). The number and location of recombinant regions was also estimated for the Taiwan-only and the PD-I/Taiwan dataset. In both instances, recombinant segments were also removed. The location and origin of recombinant segments in the Taiwan-only dataset were visualized using fastGEAR. In addition, the presence of homologous recombination between the Taiwan-only dataset and *X. taiwanensis* was evaluated following the same protocol.

### Population genetic analyses, haplotype network assessment, and population structure

Global measures of genetic diversity were estimated using the R package ‘PopGenome’ (Pfeifer et al. 2014). Briefly, the number of Single Nucleotide Polymorphisms (SNPs), nucleotide diversity (π), Tajima’s D (Tajima 1989), and the Watterson’s estimator (θ) (Watterson 1975) were computed. Nucleotide diversity (π) measures the average number of nucleotide differences per site in pairwise comparisons among DNA sequences. Tajima’s D measures the frequency of polymorphisms present in a population and compares that value to the expectation under neutrality. The Watterson θ estimator measures the mutation rate of a population. All analyses were performed using core genome alignments for the worldwide PD-causing, PD-I/Taiwan, and Taiwan-only datasets, and then repeated using their corresponding core genome alignments with removed recombinant segments (non-recombinant cores). For each dataset, isolates were divided based on their geographic origin. Tajima’s D could not be calculated from locations with small sample size (i.e., Spain, which includes only two isolates).

Haplotype networks were built for: a) the non-recombinant PD-I/Taiwan dataset, b) the recombinant Taiwan-only dataset, and c) the non-recombinant Taiwan-only dataset. Core genome haplotypes were calculated based on the number of mutations among the analyzed isolates. The haplotype network was built using the HaploNet function in the R package ‘pegas’ (Paradis 2010). Haplotypes shapes and colors were coded by location. In addition, SNP-sites (Page et al. 2016) was used to build a SNPs alignment from the non-recombinant core genome alignment for worldwide PD-causing isolates. This alignment was then used to evaluate population structure with the R package ‘hierBAPS’ (Tonkin-Hill et al. 2018).

## Results

The phylogenetic analyses (Figure 1 and Figure S1) demonstrated that all Taiwanese isolates and the PD-I lineages form a monophyletic clade, indicating a single source of introduction from the Southeast US into Taiwan. Though isolates from the same populations tended to cluster together on the phylogenetic tree, low branch support and short branch length within the PD-I/Taiwan group suggest very little sequence differentiation. This result is consistent with population structure analyses using a non-recombinant core SNPs alignment (Figure S2). However, it should be noted that the population structure analysis classified GV215, GV216, GV230, GV235, GV240, and GV249 as a distinct population from the rest of Taiwan isolates. Likewise, comparatively longer branch lengths among GV219, GV229, GV230, and GV241 were observed in the phylogenetic tree. Moreover, these four isolates formed a monophyletic clade with isolate 14B4 from Georgia. Contrary to the phylogenetic tree, the haplotype network showed two clearly differentiated groups between the Southeast US (PD-I) and Taiwan populations (Figure 2a).

**Figure 2.**
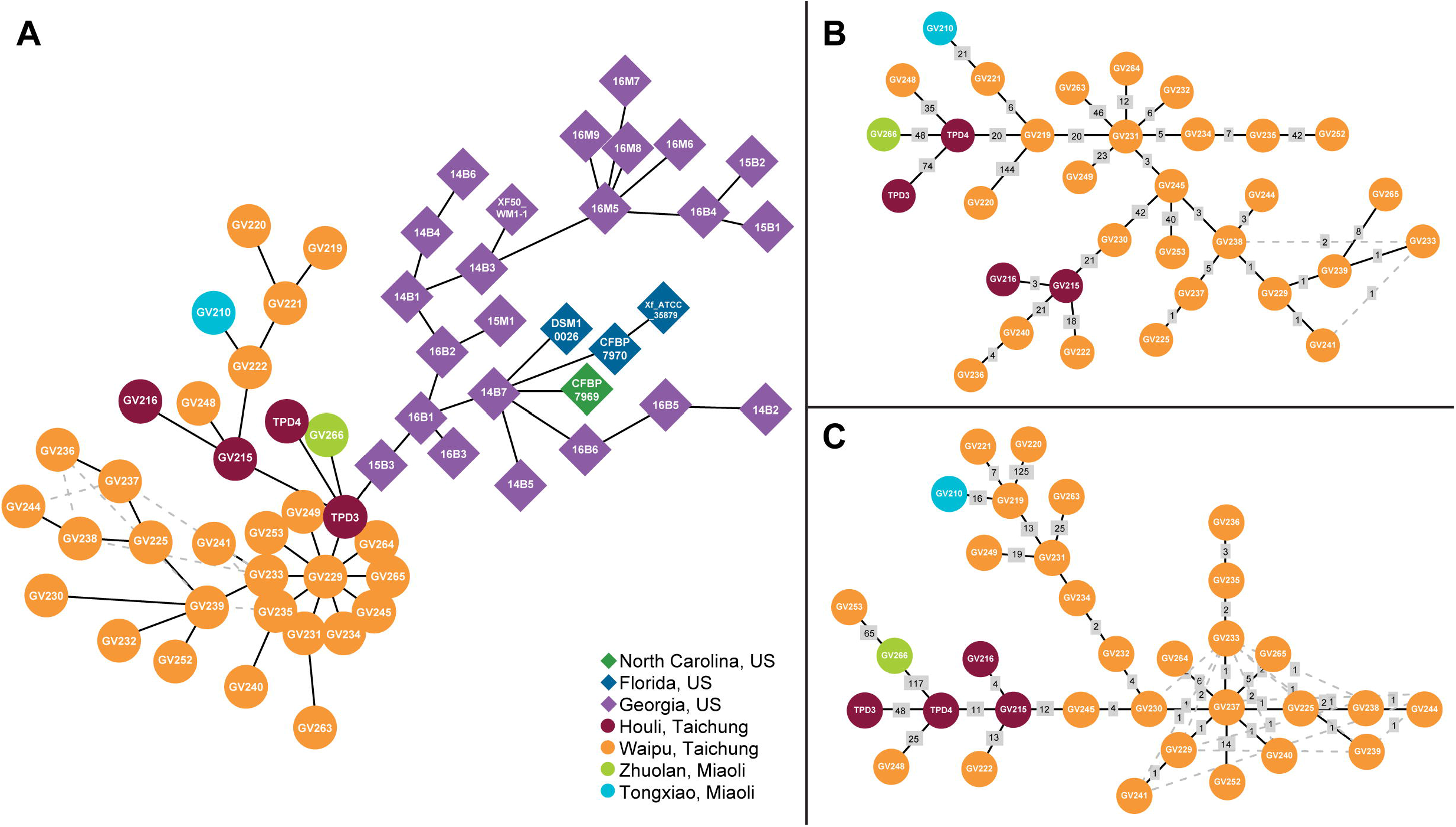
Haplotype networks showing relationships among PD-causing isolates. Isolate ID is included inside each circle/diamond. **a**. Haplotype network between Taiwan (descendant population) and PD-I/Southeast US (ancestral population) isolates. Network was built based on the core genome alignment following removal of recombinant segments of the PD-I/Taiwan dataset. **b**. Haplotype network of Taiwan isolates. Network was built based on the core genome alignment of the Taiwan-only dataset. **c**. Non-recombinant haplotype network of Taiwan isolates. Network was built based on the core genome alignment following removal of recombinant segments of the Taiwan-only dataset. Number of mutations between isolates are shown in gray boxes. Minimum spanning tree is shown in solid black lines. Alternative relationships among isolates are shown as dashed gray lines.

The core genome (i.e., shared by > 99% of strains) of worldwide PD-causing subsp. *fastidiosa* isolates included 1,715 genes, while the soft-core genome (95-99% of strains) included 258 genes. By comparison, the core and soft-core genomes in the Taiwan-only dataset included 2,041 and 67 genes, respectively (Table 2). No genes were found to be uniquely gained/loss in all Taiwan isolates in respect to the Southeast US (i.e., PD-I). A core genome haplotype network for the Taiwan-only population (Figure 2b) showed that several mutations have accumulated within the region without a clear geographic pattern. The non-recombinant core genome haplotype network confirms this trend (Figure 2c), but it also suggests that many of these mutations are likely a product of recombinant events.

**Table 2.** Summary of pan-genome analysis of worldwide PD-causing *X. fastidiosa* data and the Taiwanese population alone. Threshold for inclusion indicates the proportion of isolates harboring the genes.

| Threshold for inclusion | Worldwide (N = 206) | Taiwan-only (N = 33) |
| --- | --- | --- |
| Core (> 99%) | 1,715 | 2,041 |
| Soft-core (95-99%) | 258 | 67 |
| Shell (15-95%) | 652 | 272 |
| Cloud (< 15%) | 7,379 | 757 |

The genetic diversity between Southeast US (PD-I) (336 SNPs, π = 2.645×10e^-05^) and Taiwan (362 SNPs, π = 3.045×10e^-05^) was comparable. Likewise, the mutation rate was similar between these geographic areas (PD-I or Southeast US’s θ = 4.986×10e^-05^ and Taiwan’s θ = 5.101×10e^-05^). The similarities remained following removal of recombinant segments from the PD-I/Taiwan dataset (Table 3). The negative Tajima’s D values detected are indicative of a recent population introduction; however, values were less negative in Taiwan (−1.547) compared to the Southeast US (PD-I) (−1.857). This trend was flipped when recombinant segments were removed (−2.368 for PD-I or Southeast US and −2.717 for Taiwan).

**Table 3.** Genome-wide genetic diversity measurements.

| With recombination |  |  |  |  |  |  |
| --- | --- | --- | --- | --- | --- | --- |
| Analysis | Population (N) | Alignment length | SNPs | Pi | Watterson | Tajima |
| PD-causing isolates +<br>Costa Rica | California, USA (139) | 1,562,792 | 537 | 2.854x10e<sup>-05</sup> | 6.238x10e<sup>-05</sup> | -1.790 |
|  | Southeast USA (32) |  | 926 | 1.346x10e<sup>-04</sup> | 1.471 x10e<sup>-04</sup> | -0.329 |
|  | Taiwan (33) |  | 215 | 1.056x10e<sup>-05</sup> | 3.390x10e<sup>-05</sup> | -2.629 |
|  | Mallorca, Spain (2) |  | 2 | 1.280x10e<sup>-06</sup> | 1.280x10e<sup>-06</sup> | * |
| PD-I/Taiwan | Southeast USA (28) | 1,748,449 | 336 | 2.645x10e<sup>-05</sup> | 4.986x10e<sup>-05</sup> | -1.857 |
|  | Taiwan (33) |  | 362 | 3.045x10e<sup>-05</sup> | 5.101x10e<sup>-05</sup> | -1.547 |
| Taiwan-only | Miaoli (2) | 1,842,805 | 44 | 2.388x10e<sup>-05</sup> | 2.388x10e<sup>-05</sup> | * |
|  | Taichung (31) |  | 379 | 2.202x10e<sup>-05</sup> | 5.148x10e<sup>-05</sup> | -2.217 |
| Taiwan-only | Houli (4) | 1,842,805 | 110 | 3.364x10e<sup>-05</sup> | 3.256x10e<sup>-05</sup> | 0.349 |
| (within Taichung) | Waipu (27) |  | 312 | 1.946x10e<sup>-05</sup> | 4.393x10e<sup>-05</sup> | -2.202 |
| Without recombination |  |  |  |  |  |  |
| PD-causing isolates +<br>Costa Rica | California, USA (139) | 1,410,222 | 508 | 2.995x10e<sup>-05</sup> | 6.540x10e<sup>-05</sup> | -1.788 |
|  | Southeast USA (32) |  | 864 | 1.331x10e<sup>-04</sup> | 1.521x10e<sup>-04</sup> | -0.485 |
|  | Taiwan (33) |  | 193 | 1.044x10e<sup>-05</sup> | 3.372x10e<sup>-05</sup> | -2.633 |
|  | Mallorca, Spain (2) |  | 2 | 1.418x10e<sup>-06</sup> | 1.418x10e<sup>-06</sup> | * |
| PD-I/Taiwan | Southeast USA (28) | 1,624,189 | 272 | 1.739x10e<sup>-05</sup> | 4.345x10e<sup>-05</sup> | -2.3676 |
|  | Taiwan (33) |  | 398 | 1.767x10e<sup>-05</sup> | 6.038x10e<sup>-05</sup> | -2.7167 |
| Taiwan-only | Miaoli (2) | 1,718,980 | 119 | 6.923x10e<sup>-05</sup> | 6.923x10e<sup>-05</sup> | * |
|  | Taichung (31) |  | 361 | 1.584x10e<sup>-05</sup> | 5.257x10e<sup>-05</sup> | -2.7062 |
| Taiwan-only | Houli (4) | 1,718,980 | 51 | 1.513x10e<sup>-05</sup> | 1.618x10e<sup>-05</sup> | -0.6804 |
| (within Taichung) | Waipu (27) |  | 313 | 1.536x10e<sup>-05</sup> | 4.724x10e<sup>-05</sup> | -2.6679 |

In the Taiwan-only dataset, genetic diversity was comparable between Miaoli (44 SNPs, π = 2.388×10e^-05^) and Taichung (379 SNPs, π = 2.202×10e^-05^). On the other hand, Watterson’s θ was lower in Miaoli (θ = 2.388×10e^-05^) compared to Taichung (θ = 5.148×10e^-05^). It should be noted however that only two isolates originate from Miaoli. Within Taichung (N = 31), the genetic diversity was higher in Houli (110 SNPs, π = 3.364×10e^-05^) compared to Waipu (312 SNPs, π = 1.946×10e^-05^). Watterson’s θ showed the opposite trend (Houli’s θ = 3.256×10e^-05^ vs. Waipu’s θ = 4.393×10e^-05^). After recombinant segments were removed, genetic diversity was comparable between these two districts (51 SNPs, π = 1.513×10e^-05^ for Houli and 313 SNPs, π = 1.536×10e^-05^ for Waipu). The Watterson’s θ estimates of the non-recombinant core genome alignment maintained the same trends as those seen in the entire core genome alignment (Houli’s θ = 1.618×10e^-05^ and Waipu’s θ = 4.724×10e^-05^). Finally, Tajima’s D estimates for both districts dropped following removal of recombinant segments; however, the change was more notable in Houli (0.349 to −0.680).

Recent recombination events were observed among Taiwan isolates (Figure 3); however, they were not pervasive. Based on the fastGEAR analysis, there was no clear geographic structuring of Taiwan isolates. The recombinant segments detected originated from an unknown donor, suggesting that not all sequence diversity found within the region has been sampled. Furthermore, no core genome recombinant events were detected between the Taiwan-only subsp. *fastidiosa* dataset and *X. taiwanensis*.

**Figure 3.**
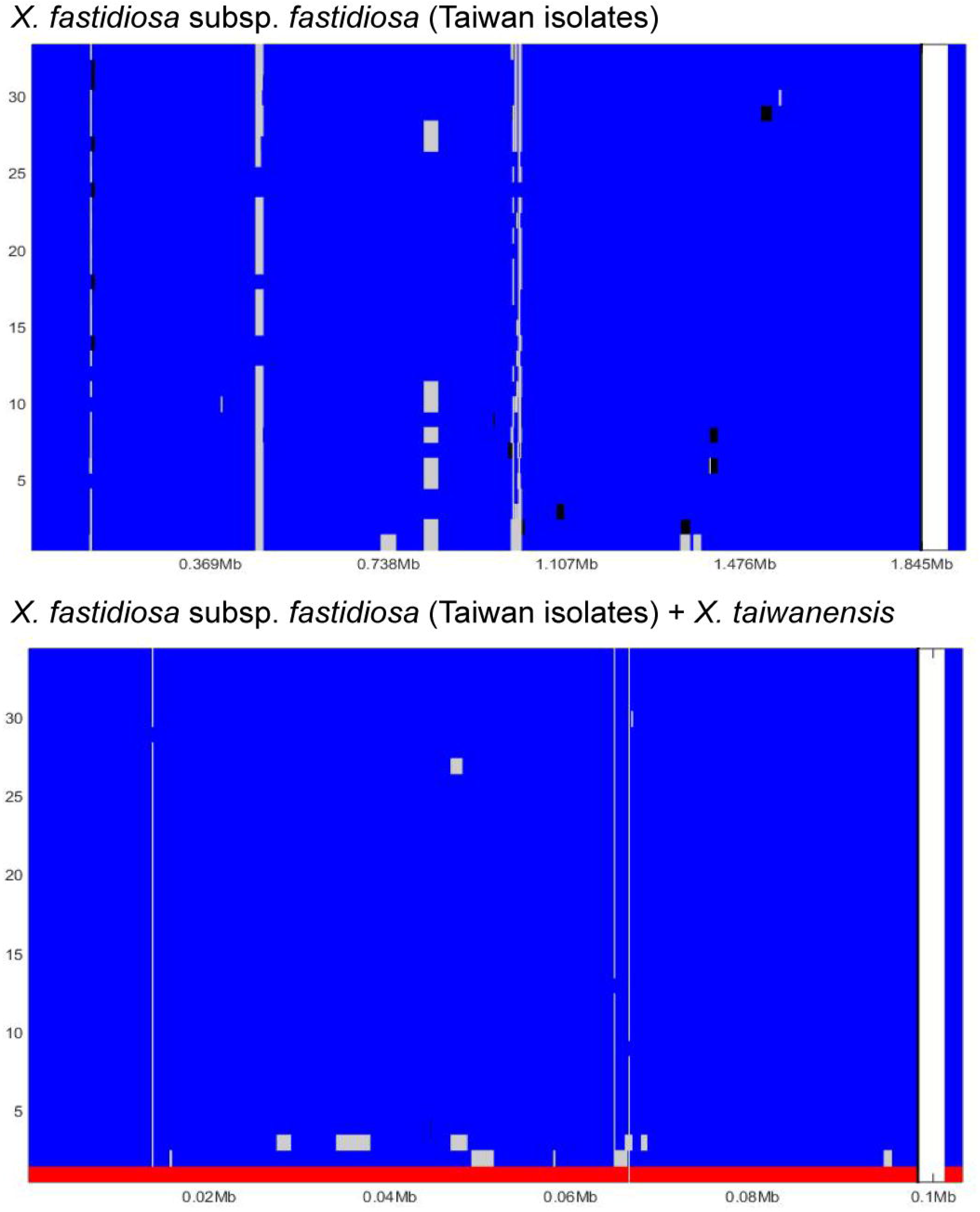
Frequency and location of recombination events in fastGEAR identified lineages. Recombination events are shown across the length of the core genome alignment of the Taiwan-only dataset. Larger areas represent recipient sequences, while shorter segments of different color within those areas represent donor sequences from another lineage. Recombinant segments from unidentified lineages are shown in black. A single lineage for all subsp. *fastidiosa* isolates in Taiwan (blue) was found. *X. taiwanensis* was identified as a second lineage (red).

## Discussion

### PD-causing subsp. *fastidiosa* was introduced into Taiwan from the Southeast US

Despite attempts at disease surveillance, quarantine, and control, novel plant pathogen outbreaks are still observed in numerous locations worldwide (Parnell et al. 2017). Following introduction, the reconstruction of invasion routes and the identification of a pathogen’s establishment and spread mechanisms becomes crucial for developing mitigation strategies (Robert et al. 2012). Here, we have confirmed previous studies suggesting that PD-causing subsp. *fastidiosa* was introduced to Taiwan from the Southeast US (Castillo et al. 2019, 2021) using multiple lines of evidence based on whole genome sequence data. The results support that the ancestor/descendant relationship between these populations was not an artifact caused by limited sampling of only two isolates from Houli in the previous study (Castillo et al. 2021).

Phylogenetic analysis demonstrated that Taiwan and Southeast US (PD-I) isolates cluster within the same monophyletic group. Moreover, short branch length and lack of phylogenetically supported geographic clusters are indicative of few nucleotide changes differentially accumulating in both populations. This finding is consistent with reports of subsp. *pauca*’s introduction into the Apulian region in Italy (Sicard et al. 2018; Saponari et al. 2018; Giampetruzzi et al. 2017). Likewise, nucleotide diversity and mutation rate were comparable between Southeast US (PD-I) and Taiwan isolates. Previous studies have found that introduced populations of subsp. *pauca* and subsp. *fastidiosa* have lower genetic diversity than native ones (Castillo et al. 2020). Yet, with time, nucleotide differences accumulate as a product of ecological and environmentally mediated pressures as well as the result of non-adaptive evolution (Castillo et al. 2021). This has been observed in the introduction of subsp. *fastidiosa* into the US ∼150 years ago (Vanhove et al. 2020). Therefore, the lack of genome-wide nucleotide differences and the similar genetic diversity and mutation rate between Taiwan and the Southeast US (PD-I) are indicative that this was a recent introduction event.

A previous study conducted using only two isolates from Taiwan found four genes gained and five loss between the ancestral Southeast US (PD-I) population and the newly introduced Taiwan population (Castillo et al. 2021). Our pangenome analysis used 33 isolates from Taiwan but failed to find a gene gain/loss pattern that could be linked to this introduction. This suggests that the previous observation was an artifact of the small sample size and that no clear gene gain/loss trends can be currently attributed to a founder effect. Furthermore, since gene gain/loss often precedes nucleotide substitutions and indels in bacterial genome evolution (Iranzo et al. 2019), this is consistent with the record of invasion of *X. fastidiosa* to Taiwan in 2002.

From the current data, it is difficult to determine if a single or multiple introduction events have taken place. In the case of subsp. *multiplex*, both phylogenetic and genetic diversity data support the hypothesis of multiple introductions into Europe from the US (Landa et al. 2020). Here, the haplotype network analysis is indicative of a single introduction. However, it is possible that genetic diversity losses associated with a founder event could mask the existence of simultaneously introduced strains. The population structure analysis shows evidence of genetic differentiation of two strains from Houli (GV215 and GV216), and four strains from Waipu (GV230, GV235, GV240, and GV249) into a distinct group. Yet, these strains are neither phylogenetically nor geographically aggregated. Whether the distinction of these strains as a genetically unique population in Taiwan is the result of accumulated nucleotide substitutions or the product of a distinct introduction event remains to be determined.

### Taiwan isolates are quickly differentiating via multiple mechanisms

It should be noted that despite evidence suggesting a short evolutionary time since the introduction of PD-causing subsp. *fastidiosa* to Taiwan, sequence divergence in core genome genes is already observed in the region. It is not possible to confidently ascertain if this is due to variable evolutionary pressures across locations in Taiwan or an intrinsic genetic characteristic carried over from the ancestor population. Waipu and Houli are neighboring districts that are < 10 km apart and share similar warm humid subtropical climate conditions, so it would be expected that environmental pressures would be similar among populations. Likewise, Zhuolan and Tongxiao are separated from Taichung City by only ∼20 km. This proximity could also explain the lack of clear phylogenetic differentiation among isolates from distinct locations. However, while isolates obtained from infected grapevines in either region could not be phylogenetically separated, trends in genetic diversity and mutation rates did vary among locations. Genetic diversity was comparable between Miaoli and Taichung despite the differences in sample size. Among Taichung locations, genetic diversity in Houli was higher than that observed in Waipu, so it is possible that certain biotic or abiotic conditions do favor the accumulation of mutations in this location. Nonetheless, future studies should evaluate if the observed differences are simply due to low sample sizes from regions outside of Waipu.

Another notable aspect is how global diversity estimates in Houli and Waipu responded to the removal of recombinant regions in the core genome alignment. Further supporting this observation, are the changes in the minimum spanning tree seen on the Taiwan-only haplotype network following recombination removal. When recombinant segments were included, the minimum spanning tree (solid black lines) was defined and showed no clear separation among Taiwan isolates from different districts or townships (Figure 2b). Following the removal of recombinant segments, Houli isolates clustered within the network and the relationships among Waipu isolates became less clear (dashed gray lines) (Figure 2c). This is evidence that recombination significantly contributes to the genetic differentiation of PD-causing subsp. *fastidiosa* in Taiwan.

However, the changes in global diversity seen following the removal of recombinant events were more dramatic in Houli compared to Waipu. Previous studies have shown that natural competency and recombinant capacity are isolate dependent (Vanhove et al. 2019; Nunney et al. 2012, 2014; Landa et al. 2020). Moreover, recombinant events can be carried over from ancestral populations into new introductions; particularly, in instances where these introductions are recent (Landa et al. 2020). Therefore, our results are indicative that recombinant-prone subsp. *fastidiosa* isolates might have been introduced to Houli from the Southeast US or that they might have evolved there. Further sampling within this region would be required to evaluate this. Nonetheless, the existence of unknown recombining sequences in both districts suggest that genetic diversity within Taiwan is higher than what can be defined via our current sampling. Finally, even though homologous recombination is known to occur in sympatric subsp. *pauca* populations (Almeida et al. 2008) and among different *X. fastidiosa* subspecies (Nunney et al. 2014), no recombination events between PD-causing subsp. *fastidiosa* and *X. taiwanensis* were detected. This result is expected based on the low levels of sequence identities between these two species at ∼84% (Su et al. 2016; Weng et al. 2021) and further support that these two species are ecologically and/or biologically isolated.

## Conclusion

In globally distributed pathogens, understanding the relationships among populations and the location-specific mechanisms of diversification has significant implications in the development of successful management techniques. Here, we used phylogenetic analysis to investigate the introduction of subsp. *fastidiosa* into Taiwan and confirm that the invasion originated from a single source population (i.e., the Southeast US). Haplotype network analysis, pan-genome trends, and global genetic diversity measures support this hypothesis. Following its recent establishment within Taiwan, PD-causing subsp. *fastidiosa* has differentiated across geographic locations. We propose that part of this diversification can be linked to homologous recombination acting more predominantly within Houli, Taichung.

## Data Statement

All raw reads have been deposited in NCBI under BioProject PRJNA715299. The complete genome sequence of the reference isolate GV230 has been deposited in GenBank/ENA/DDBJ under the accession CP060159.

## Funding

Funding was provided by the Ministry of Science and Technology of Taiwan (MOST 106-2923-B-002-005-MY3) to CWT and Academia Sinica to CHK. The work also received funding from the PD/GWSS Research Program, California Department of Food and Agriculture, and the European Union’s Horizon 2020 research and innovation program under grant agreement no. 727987, “*Xylella fastidiosa* Active Containment Through a multidisciplinary-Oriented Research Strategy XF-ACTORS”. The funders had no role in study design, data collection and interpretation, or the decision to submit the work for publication.

## Acknowledgements

The Illumina and Oxford Nanopore sequencing library preparation services were provided by the Genomic Technology Core (Institute of Plant and Microbial Biology, Academia Sinica). The Illumina MiSeq sequencing service was provided by the Genomics Core (Institute of Molecular Biology, Academia Sinica). We thank Yi-Ming Tsai for technical assistance.

## Conflict of Interest

The authors declare no conflict of interest.

## Supplementary Materials

**Figure S1. Maximum Likelihood (ML) non-recombinant tree of worldwide PD-causing infecting subsp. *fastidiosa* isolates**. The phylogeny was constructed using the core genome excluding the recombinant segments detected by fastGEAR. Costa Rican isolates are used to root the tree. Clades encompassing California and Spain isolates have been compressed to their most recent common ancestor.

**Figure S2. Evolutionary relationship of populations identified by hierBAPS**. Non-recombinant ML cladogram showing the clustering of isolates from hierBAPS identified populations at level 2. Clustering analysis was performed using a non-recombinant SNP alignment. Different populations are represented by distinct colored dots at branch tips. Major geographical populations are separated by vertical lines.

## Notes

### Competing Interest Statement

The authors have declared no competing interest.

