## Supplementary figures and images for "Genetic differentiation of *Xylella fastidiosa* following the introduction into Taiwan"

### Figure S1

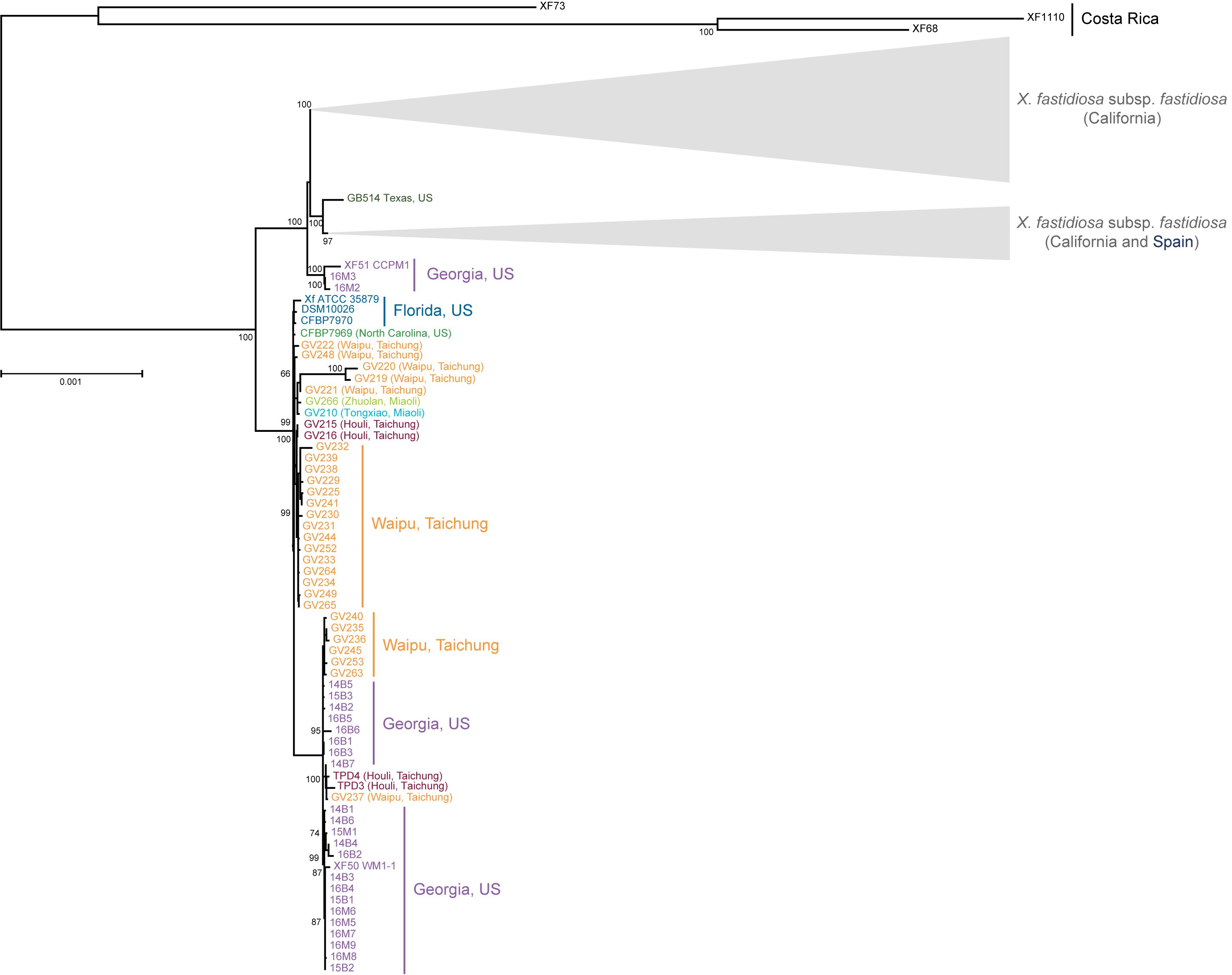

### Figure S2

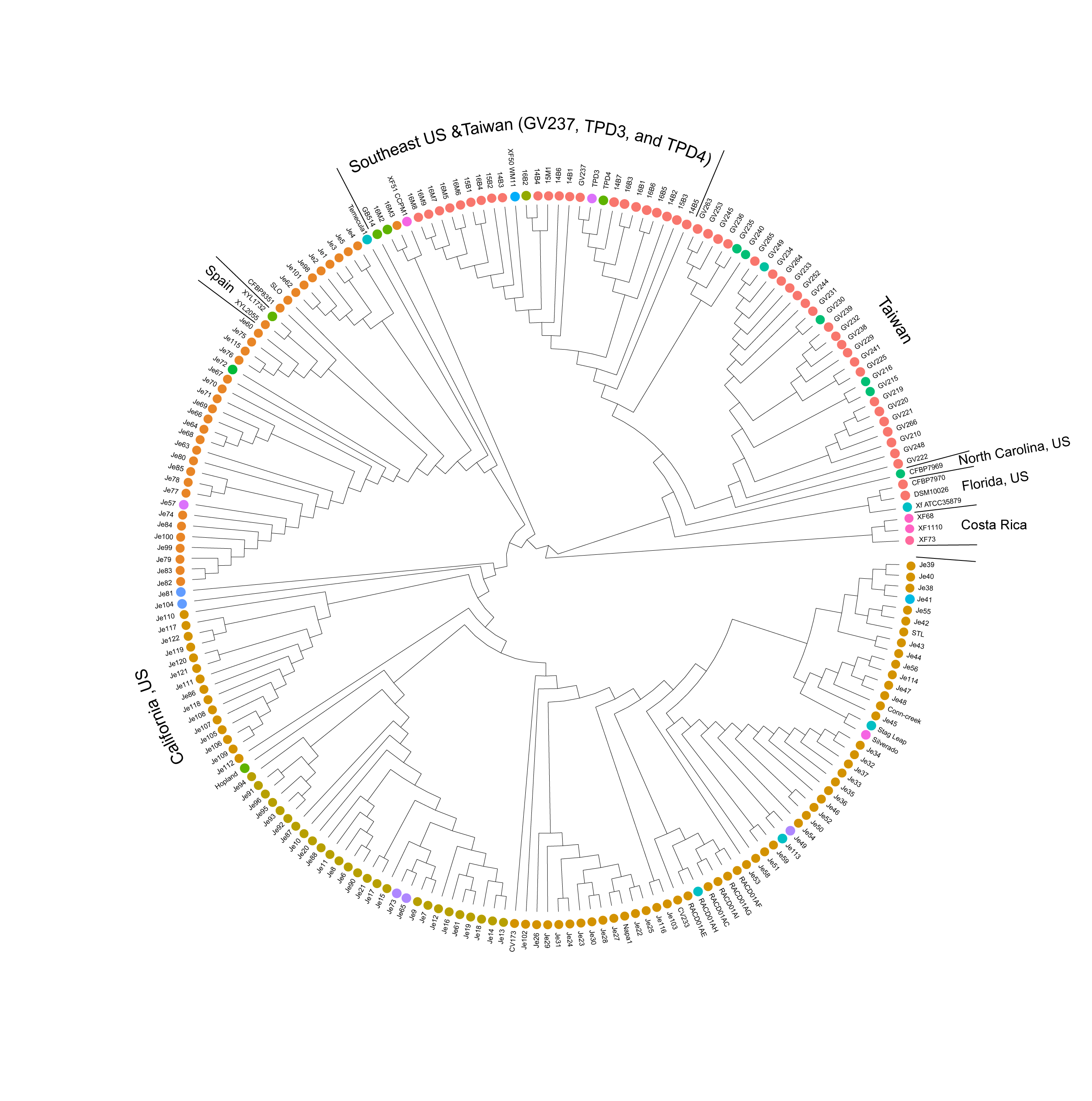
